# Allelic variations and gene cluster modularity act as non-linear bottlenecks for cholera emergence

**DOI:** 10.1101/2022.09.26.509565

**Authors:** Mario López-Pérez, Deepak Balasubramanian, Cole Crist, Trudy-Ann Grant, Jose M. Haro-Moreno, Asier Zaragoza-Solas, Salvador Almagro-Moreno

**Author notes:** Address correspondence to: Salvador Almagro-Moreno. Contributed equally (Mario López-Pérez, Deepak Balasubramanian).

## Abstract

Cholera, an acute secretory diarrhea, is caused by strains from a phylogenetically confined group within the *Vibrio cholerae* species, the pandemic cholera group (PCG). To date, the molecular and evolutionary factors that enable the isolated emergence of toxigenic *V. cholerae* from environmental populations remain mostly enigmatic. Comprehensive analyses of over 1,100 *V. cholerae* genomes, including novel environmental isolates from this study, reveal that the species consists of four major clades and several minor ones. PCG belongs to a large clade located within a lineage shared with environmental strains, the pandemic cholera lineage. This hierarchical classification provided us with a framework to unravel the eco-evolutionary dynamics of the genetic determinants associated with the emergence of toxigenic *V. cholerae*. Our analyses indicate that this phenomenon is largely dependent on the acquisition of unique modular gene clusters and allelic variations that confer a competitive advantage during intestinal colonization. We determined that certain PCG-associated alleles are essential for successful colonization whereas others provide a non-linear competitive advantage, acting as a critical bottleneck that elucidates the isolated emergence of PCG. For instance, toxigenic strains encoding non-PCG alleles of a) *tcpF* or b) a sextuple allelic exchange mutant for genes *tcpA*, *toxT*, *VC0176*, *VC1791*, *rfbT* and *ompU*, lose their ability to colonize the intestine. Interestingly, these alleles do not play a role in the colonization of model environmental reservoirs. Our study uncovers the evolutionary roots of toxigenic *V. cholerae* and offers a tractable approach for investigating the emergence of pathogenic clones within an environmental population.

**SIGNIFICANCE:** The underlying factors that lead to specific strains within a species to emerge as human pathogens remain mostly enigmatic. Toxigenic clones of the cholera agent, *Vibrio cholerae*, are encompassed within one phylogenomic clade, the pandemic cholera group (PCG). Here, we investigate the molecular and evolutionary factors that explain the confined nature of this group. Our analyses determined that the emergence of PCG is largely dependent on the acquisition of unique modular gene clusters and allelic variations that confer a competitive advantage during intestinal colonization. These allelic variations act as a critical bottleneck that elucidates the isolated emergence of PCG and provides a tractable blueprint for the study of the emergence of pathogenic clones within an environmental population.

## INTRODUCTION

The emergence of human pathogens is one of the most pressing public health concerns of modern times, with new outbreaks of varying magnitude occurring regularly around the world^1–4^. Pathogen emergence is a complex and multifactorial phenomenon that culminates with a microorganism acquiring the ability to colonize and harm the human host, cause outbreaks, and persist within the environment^5^. To date, the specific factors and evolutionary constraints that govern the emergence of bacterial pathogens from environmental populations remain mostly enigmatic^4^. Elucidating how the interplay between the numerous molecular and environmental drivers lead to this complex phenomenon is critical to develop surveillance platforms to predict emergent events and potential sources of outbreaks^4^. Furthermore, in addition to the profound public health implications, understanding pathogen emergence addresses one of the central questions of microbial ecology: what are the evolutionary processes within a bacterial population that underlie colonization and adaptation to a new environment?^6^.

The Vibrionaceae is a highly diverse family of aquatic bacteria that encompass several species representing distinct paradigms of how pathogens can emerge from environmental populations^7,8^. *Vibrio cholerae* is the most widely studied member of the Vibrionaceae and has been a model organism for numerous molecular, ecological, and epidemiological studies^9–12^. Toxigenic *V. cholerae* is the causative agent of the severe diarrheal disease cholera, a scourge that affects millions of people each year and is responsible for over 100,000 deaths annually^9^. The first six pandemics of cholera are thought to have been caused by the Classical biotype *V. cholerae* O1, a strain that is now considered extinct^9,13–15^. The seventh and current pandemic is caused by the El Tor biotype of *V. cholerae* O1 and started in Indonesia in 1961^16,17^. The disease typically affects nations with limited access to clean water and poor sanitation and can be lethal if not properly treated^9,14^. Interestingly, only a limited number of *V. cholerae* strains can cause pandemic cholera^18–20^. Specifically, toxigenic strains of *V. cholerae* are confined to a phylogenetic group, what we term the pandemic cholera group (PCG)^19,21–23^. Even though the mechanisms of cholera pathogenesis in humans have been extensively studied over the last several decades, the evolutionary events and molecular constraints that explain why only members from PCG can cause pandemic cholera remains unknown^24–29^.

Some of the major general events that are essential for the emergence of toxigenic *V. cholerae* from environmental populations include the acquisition of the lipopolysaccharide (LPS) O1 antigen cluster and mobile genetic elements (MGEs) such as CTXΦ phage, Vibrio pathogenicity Island-1 (VPI-1) and Vibrio pathogenicity Island-2 (VPI-2)^15^. Among several other differences with Classical, El Tor strains have also acquired two unique pathogenicity islands (PAIs): *Vibrio* Seventh Pandemic (VSP) I and II. CTXΦ encodes the genes for the cholera toxin, which is responsible for the severity of the profuse diarrhea associated with cholera^30^. The toxin coregulated pilus (TCP), an essential colonization factor, is encoded within VPI-1^31,32^, which also encodes several major virulence regulators such as ToxT and TcpP^32,33^. VPI-2 encodes the genes for sialic acid utilization, which confers a competitive advantage in the gut^34^. VSP-I codes for a regulator that plays a role in intestinal colonization and chemotaxis^35^ and VSP-II is associated with environmental survival and fitness of the bacterium^36–38^. It was recently found that VSP-II and VPI-2 encode systems that prevent the uptake of foreign genetic material^39^. These MGEs are always found in the PCG but are not exclusive to them as they are also encoded in some environmental strains of *V. cholerae*^40–42^. However, to date, their abundance, specific distribution, and dynamics within the *V. cholerae* species are not well understood.

Horizontal acquisition of key virulence genes is a critical step in the emergence of toxigenic clones of *V. cholerae*, however, it is not sufficient to explain why the ability to cause cholera is limited to strains belonging to the PCG. This phylogenetically confined distribution together with the presence of CTXΦ and the PAIs in some environmental strains strongly indicate that other barriers and requirements exist that stringently limit their emergence. We recently investigated the genomic prerequisites that must be present in a population before a pathogenic clone can emerge from an environmental gene pool^22^. We determined that the environmental ancestor of the PCG had a particular genomic background containing a specific set of alleles of core genes that conferred preadaptations to virulence and enhanced its pathogenic potential, what we term virulence adaptive polymorphisms (VAPs)^22^. Interestingly, our data shows that VAPs are present in environmental populations and confer virulence-associated traits to strains encoding them. Overall, our results indicate that there are core genomic factors beyond the acquisition of MGEs that are critical for the emergence of toxigenic *V. cholerae*. To date, the unique core genes, and allelic variations in the genomic background of toxigenic *V. cholerae* that explain their narrow distribution remain enigmatic.

In this study, we investigated the evolutionary constraints that govern the emergence of bacterial pathogens from environmental populations using toxigenic *V. cholerae* as a model system. To do this, first, we isolated environmental *V. cholerae* strains from the Indian River Lagoon (IRL), a highly diverse and expansive estuary located in Eastern Florida. Genome comparisons of these strains against *V. cholerae* PCG unveil a phylogeny in which strains form cluster-like groups. We extended our analyses to over 1,100 publicly available genomes and found that *V. cholerae* strains are divided into four major clusters and numerous other minor clades. We found that PCG belong to one large cluster (C24) and is located within one lineage shared with other environmental strains, the pandemic cholera lineage (PCL). Subsequently, we examined the genetic determinants associated with the emergence of toxigenic *V. cholerae* through multi-level genomic analyses by investigating **a)** the distribution and abundance of the O1 LPS cluster and main virulence associated MGEs, **b)** the evolutionary dynamics of the gene clusters that comprise these MGEs, and **c)** the allelic diversity within the population. We found that these elements are made up of several modules sparsely distributed among the different phylogenomic groups and include small gene clusters and allelic variations. We determined that these allelic variations confer a competitive advantage during intestinal colonization, acting as a critical bottleneck that elucidates the isolated emergence of PCG, but do not play a role in the colonization of model environmental reservoirs. Overall, our study provides a multi-level scenario that serves as a blueprint for the study of the evolution of facultative bacterial pathogens.

## RESULTS AND DISCUSSION

### Isolation of environmental *Vibrio cholerae* from the Indian River Lagoon

*V. cholerae* is a natural inhabitant of aquatic ecosystems where it can be found as free-living or attached to abiotic or biotic surfaces^43,44^. We recently established two sampling sites along the eastern coast of Florida where we isolated pathogenic strains of *Vibrio vulnificus*^45^. We sampled those two sites, Feller’s House (FH) and Shepard Park (SP), to investigate the potential presence of *V. cholerae* strains (**Fig. 1A**). Using the isolation, enrichment and selection methods described below, we identified a total of 178 potential *V. cholerae* isolates from the different fractions at both locations during our sampling events (42 FH and 136 SP). To confirm those isolates, we performed a PCR screening for the species-specific gene *ompW*^46^. Only 65 isolates were confirmed as *bona fide V. cholerae*, all of them isolated from SP (**Fig. 1A**). *V. cholerae* isolates from SP were collected from all the fractions that we examined (sediment, cyanobacteria, particle-associated and water) except for oysters. Subsequently, we performed PCR-based fingerprinting of the strains to reduce redundancies and minimize clonality. We used a combination of PCR-based fingerprinting analyses, VCR-PCR and ERIC-PCR^47^. Based on fingerprinting patterns and cluster analysis performed using BIONUMERICS v8.0 we generated a dendrogram to group these observed patterns. Using this resulting dataset, we selected 35 isolates with unique fingerprinting patterns for whole genome sequencing (**Fig. S1 and Table S1**).

**Fig. 1.**
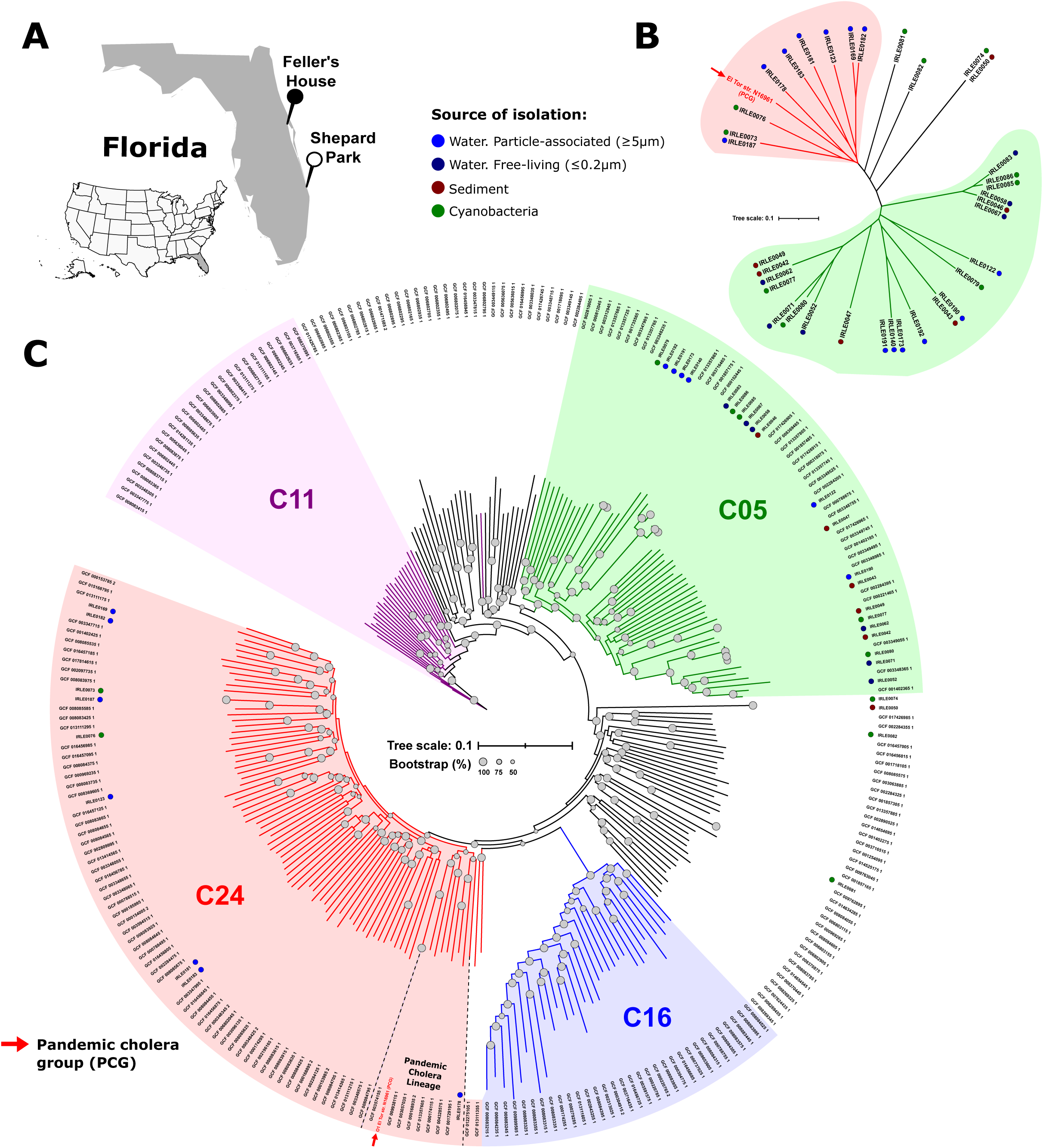
Phylogenomic and population structure of *Vibrio cholerae*. **(A)** Map of Florida indicating the locations where the environmental *V. cholerae* strains were isolated (Shepard Park and Felleŕs House). **(B)** Whole-genome SNP-based phylogenetic tree showing the relationships among the *V. cholerae* genomes obtained in this study and a representative strain from the Pandemic Cholera Group (highlighted with a red arrow). **(C)** Phylogenomic tree of dereplicated *V. cholerae* strains available in public databases and strains from this study. The tree was reconstructed from single nucleotide polymorphisms of the core genome. Branches of members belonging to the four main clusters are colored in green (C5), violet (C11), blue (C16) and red (C24). The colored circles represent the source of isolation of the environmental strains collected in this study. Grey circles located at the branch nodes represent the bootstrap values.

### Phylogenomic characterization and clustering of *V. cholerae* strains

The 35 *V. cholerae* strains that we sequenced were isolated from a diverse set of fractions: 6 from sediment, 10 from cyanobacteria, 9 from the free-living water fraction and 10 from the particle-associated one (**Table S1**). We examined the relationship of these strains with *V. cholerae* PCG by analyzing the average nucleotide identity (ANI) in a pairwise comparison against *V. cholerae* N16961, a reference O1 strain from the El Tor biotype (**Fig. S2**) ^48^. The minimum values obtained were ca. 97.5, always within the established cut-off for strains of the same species. However, up to nine environmental strains showed a greater divergence with the PCG (ANI >98.2%) than with the rest of the strains (**Fig. S2**). Interestingly, phylogenomic analysis based on whole-genome single nucleotide polymorphism (SNP) (230,824 SNPs identified in 3.19 Mb of core genome) show that the majority of the IRL strains form two major clusters, one of them including the toxigenic strain of *V. cholerae* (**Fig. 1B**). This evolutionary relationship shares similarity with that of *V. vulnificus*, in which the pathogenic strains capable of causing septicemia in humans belong to one clearly defined cluster^49^. This prompted us to further characterize the *V. cholerae* species to investigate whether the evolution of the toxigenic strains follow a similar cluster-based pattern, as this would shed important light on the emergence and evolution of the pathogen.

First, we collected the *V. cholerae* genomes available in public databases (based on NCBI classification accessed in July 2020; see Materials and Methods) together with our environmental isolates. Naturally, there is a significant bias in the databases towards *V. cholerae* O1 and O139 strains, which are mostly clonal. Therefore, in order to obtain an unbiased phylogeny, we dereplicated all these genomes to an identity of ANI >99%. From a total of 1,127 initial genomes, we obtained 227 non-clonal reference *V. cholerae* clades (**Table S2**). To maximize the consistency of our analyses, we built both a SNP-based phylogeny and genome clustering using ANI to determine the groups that constitute the *V. cholerae* species. The application of a divergence threshold of less than 97% in ANI results in a single cluster indicating that the boundary of intra-population sequence diversity in *V. cholerae* species is 97%. However, clustering using a threshold of 98% of ANI shows that the species encompasses 34 clusters that include *V. cholerae* strains (**Table S3**). Next, we used a total of 458,798 SNPs from a core genome size of 2.46 Mb to build the species phylogeny using non-clonal reference clades and IRL strains (**Fig. 1C**). The species phylogeny topology based on SNPs or ANI clustering are virtually identical, with some minor exceptions that might be due to certain strains having a convergent history. Interestingly, both approximations reveal that four major clusters account for 69.2% of the non-clonal clades (157 out of a total of 227) and 31 of the 35 IRL isolates from this study (**Table S3**). All IRL isolates are grouped together with previously isolated strains except for strains IRLE0050 and IRLE0074 which formed a separate cluster (**Table S3**). Cluster 24 (C24) is the largest of the four main clusters containing 68 non-clonal clades including PCG. On the other hand, most IRL strains belong to C5 (22 of the 35) (**Fig. 1C**). We did not find a pattern that associate the phylogeny of the IRL isolates and their fraction of isolation. The C24 phylogeny also reveals that PCG groups with nine other non-clonal clades, including IRLE0178 from this study, fall into a separate branch in the phylogenomic tree, what we have termed the Pandemic Cholera Lineage (PCL) (**Fig. 1C**). This provides us with a novel hierarchical approach based on three levels to decipher the evolutionary history of toxigenic strains of *V. cholerae* (from highest to lowest): a) the C24 within all *V. cholerae* strains, b) the PCL within C24 non-clonal clades and c) the clonal *V. cholerae* toxigenic strains within the PCL lineage.

### Lipopolysaccharide (LPS) cluster

The LPS plays numerous roles in the pathogenesis of toxigenic *V. cholerae* (e.g., endotoxic activities, immunological responses, etc.^50–52^) and is critical to determine the antigenic properties and classification of the bacterium^52^. Specifically, given that only O1 strains, together with the extinct O139, can cause pandemic cholera, we examined its LPS cluster to determine its distribution, evolutionary history, and allelic diversity within the *V. cholerae* species (**Fig. 2**). The O1 gene cluster is 29.7 Kb long and encompasses approximately 26 ORFs with potential or demonstrated function (some are truncated preventing an exact number). The *rfa* gene cluster (LPS core synthesis) is encoded at the 5’ end of the island (**Fig. 2A** blue arrows) and is widely distributed across all clades, with the notable exception of RfaF (ADP-heptose--LPS heptosyltransferase 2, VC_0235) and RfaL (O-antigen ligase, VC_0237) present in ca. 35 non-clonal clades. Nonetheless, the genes in the *rfa* cluster exhibit the highest divergence within the LPS cluster (median identity 86-88%) and accumulate numerous synonymous polymorphisms (**Fig. 2A**). This indicates that, like in *V. vulnificus*, the *rfa* cluster appears to be a preferred site to act as hot spots for the recombination of this island due to the conservation and synteny of these genes^49^. The distribution of the central region of the LPS cluster is much more restricted and is only present in 17 *V. cholerae* non-clonal clades, however, those that encode it share 100% identity with the region of PCG. This region contains the *rfb* cluster which is associated with antigen-O synthesis^53^ (**Fig. 2A**, green arrows). Interestingly, only 5 out the 17 clades that encode this region belong to the PCL, indicating that there is no direct correlation between the phylogeny and its presence in PCG. The LPS cluster of PCG encodes four transposases, which likely increase the variability within the island (**Fig. 2A and 2B**). These IS elements are widely distributed among *V. cholerae* strains providing insight into how small gene clusters can be introduced in the island. For instance, even though strains VC22 (C24) and 2479-86 (C27) encode the same version of the PCG LPS cluster (identity >99%), they do not encode two of these transposases (**Fig. 2B**). Interestingly, these 2 transposases are inserted within *rfbT* in PCG, resulting in a truncated, non-functional version of the gene (**Fig. 2B and 2C**). *rfbT* encodes a methyltransferase that is responsible for seroconversion between *V. cholerae* O1 Ogawa and Inaba serotypes, suggesting a major role for these IS sequences in the emergence of PCG^54^. Furthermore, several strains from C23 encode the same version of the hypervariable region as VC22 and 2479-86 (identity 99%) but the *rfa* cluster is different (**Fig. 2B**). The first three genes of the island encode different alleles of *rfaF*, *rfaG* and *rfaL*, and the identity of the remaining three is 95% (**Fig. 2B**). The *rfa* cluster pattern of these strains is similar to that of strain A325 (C30) (**Fig. 2B**). In addition, in this strain, the two transposases that truncate the *rfb*T gene in the PCG are inverted maintaining an intact version of the gene (**Fig. 2C**). Overall, we found a remarkable degree of conservation of the LPS cluster, which is present in almost twenty clades encoding a PCG-like version. This is particularly striking as LPS hypervariability is a universal bacterial characteristic, suggesting some strong evolutionary pressure maintaining this version of the island widespread in the population. It appears that the island is a) exchanged by homologous recombination leading to the whole gene cluster being conserved and b) the presence of IS elements increases the variability by truncating genes such as *rfbT*.

**Fig. 2.**
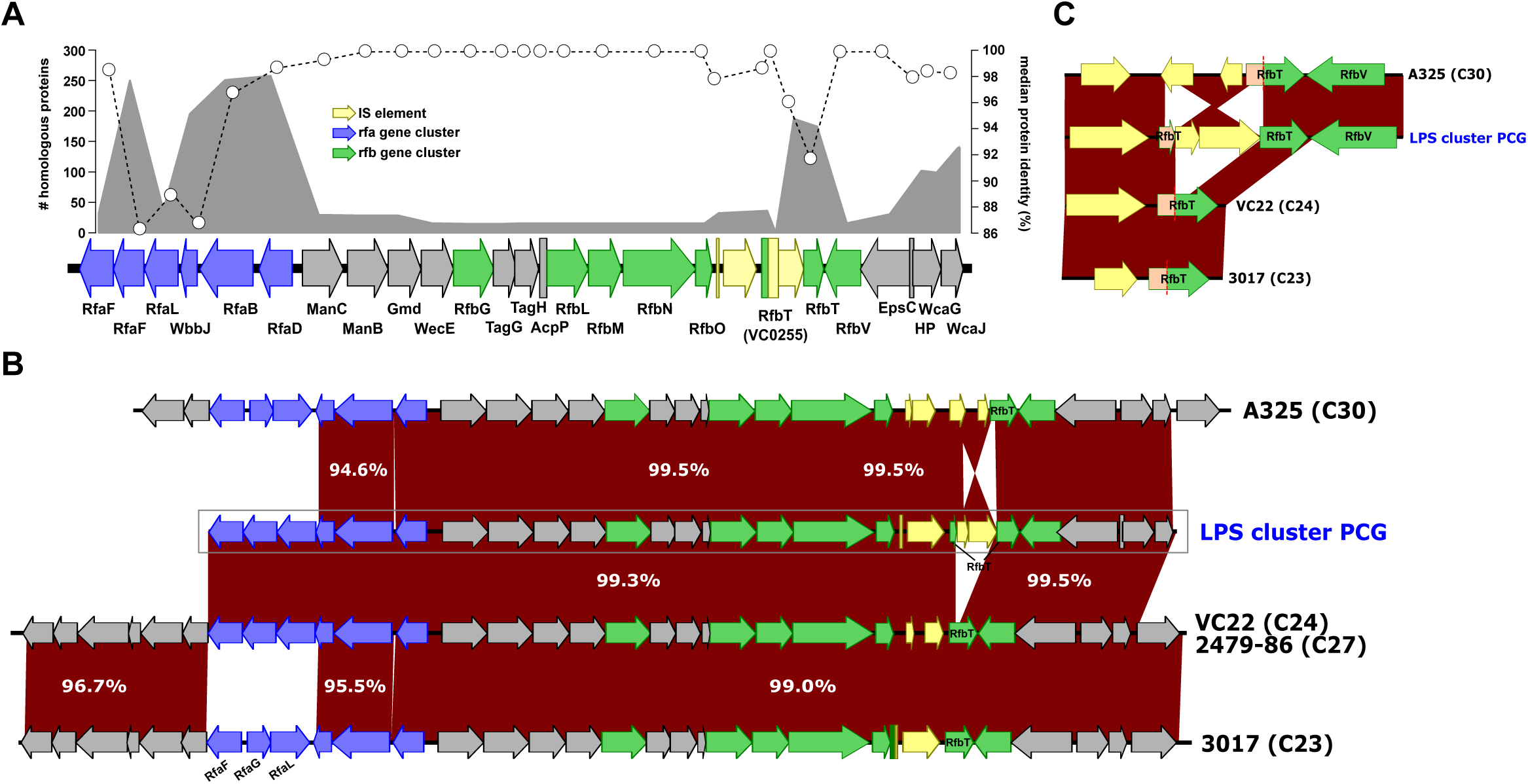
Distribution and evolutionary dynamics of the O1 Lipopolysaccharide (LPS) cluster in the *V. cholerae* species. **(A)** The area outlined in gray represents the number of homologous proteins found in the dereplicated *V. cholerae* genomes. The white circles with dotted lines indicate the median protein identity. **(B)** Genomic comparison of PCG-like LPS clusters identified in *V. cholerae* environmental genomes. **(C)** Schematic representation of the variable zone of the LPS containing the *rfb*T gene and the IS elements inserted in the PCG allele. IS elements are highlighted in yellow.

### Abundance and distribution of virulence-associated mobile genetic elements in V. cholerae species

There are several horizontally acquired MGEs that are directly associated with the emergence of toxigenic clones of *V. cholerae*. After we established the phylogeny and clustering of the *V. cholerae* species, we investigated the distribution, evolutionary dynamics, and allelic diversity of these elements within the population.

#### 1. CTXΦ phage

The genes for cholera toxin (CT) are encoded within the CTXΦ phage^30^. The production of CT in the intestine is responsible for the severity of the profuse diarrhea associated with cholera. CT leads to the secretion of electrolytes and nutrients such as fatty acids into the lumen of the small intestine^9,55,56^. Due to the lysogenic nature of CTXΦ, the insertion and deletion dynamics of this phage enables recombination of genes, which leads to diversity within the toxigenic strains ultimately serving as an opportunity to increase their pathogenic potential^30,57,58^. Inquiry into the distribution and composition of this element within *V. cholerae* shows that there is no environmental strain that recapitulates all the phage genes of PCG. At most, five clades and two clusters encode 10 of the 14 genes that make up the CTXΦ phage of *V. cholerae* N16961: three from C24 (represented by strains AAS91, PivertUAT4Aug and V060002) and two from C17 (3541-04 and 3528-08) (**Fig. 3A**). Four of these strains encode the two subunits of the cholera toxin (CtxAB). However, genes coding for the duplicated transcriptional regulator RstR (VC_1455 and VC_1464) are only found in one strain: AAS91 (**Fig. 3A**, brown shade). This strain belongs to the PCL and encodes the complete version of CTXΦ except for VC_1460 that encodes a minor coat protein pIII related to host interaction as part of the infection process (**Fig. 3B**)^59^. Instead, a transposase has been inserted in this locus in strain AAS91, followed by a APH(3’)-II family aminoglycoside O-phosphotransferase (**Fig. 3B**). In addition, AAS91 has a third copy of the triplet (RstA-RstB-RstR) instead of the two copies present in PCG strains (**Fig. 3B**). The second most complete version of CTXΦ in *V. cholerae* is found in strains V060002 from PCL and 3528-08 (C17). Both strains have lost the gene coding for RstC and encode a different allele of the transcriptional regulator RstR (**Fig. 3B**).

**Fig. 3.**
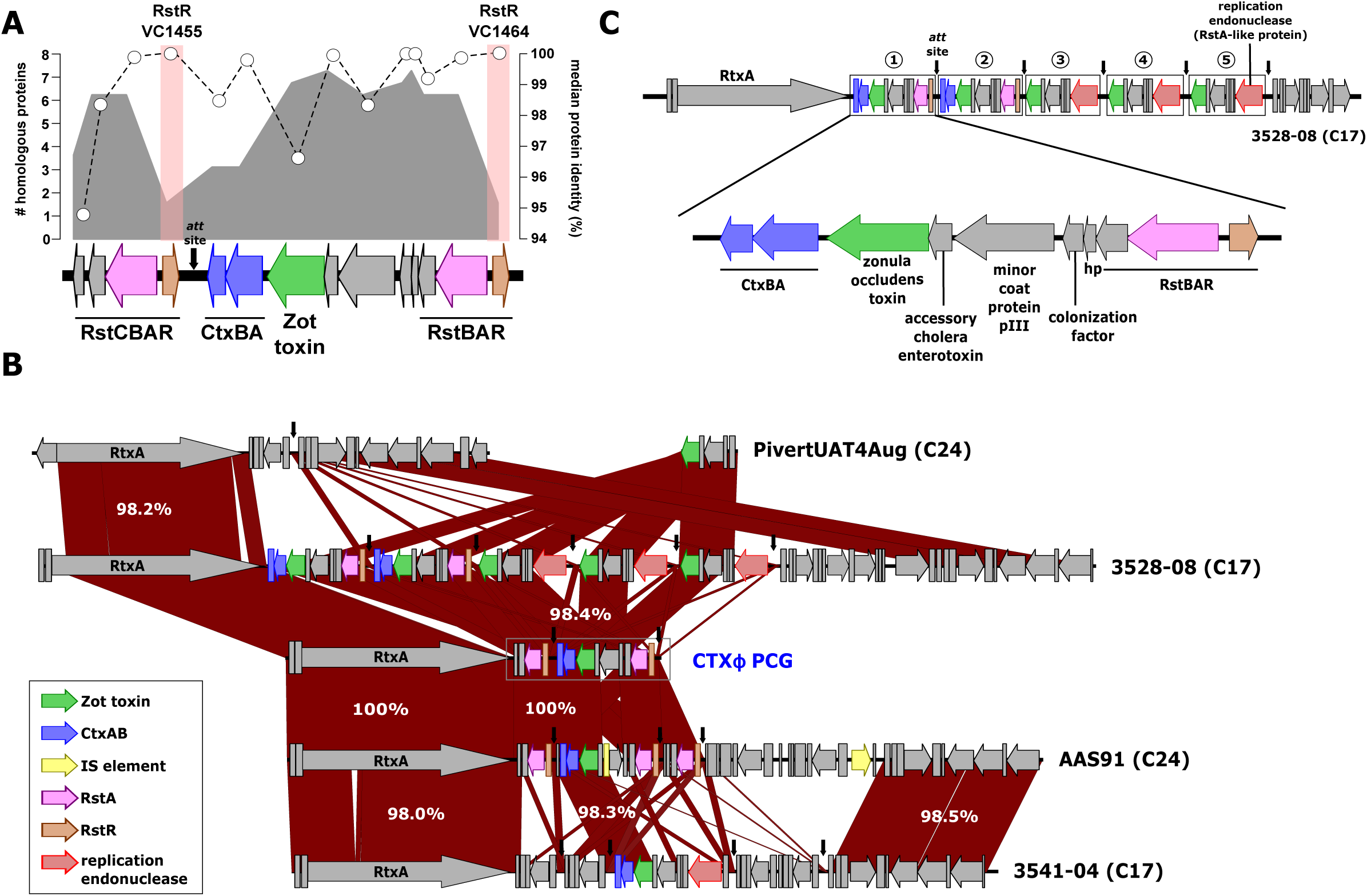
Integron-like dynamics govern genetic variation of CTXΦ phage. **(A)** Distribution and structure of the CTXΦ phage in the *V. cholerae* species. The area outlined in gray represents the number of homologous proteins found in the dereplicated *V. cholerae* genomes. The white circles with dotted lines indicate the median protein identity. Red rectangles on the genes indicate their exclusive presence in the PCL. The black arrows indicate the genomic position where the cassettes have been inserted (*att* site). **(B)** Genomic comparison of similar CTXΦ phage found in dereplicated *V. cholerae* genomes. **(C)** Schematic representation CTXΦ phage found in *V. cholerae* 3528-08, both the numbers and the rectangles mark the different clusters acquired. Clusters of genes with a known function follow the same color pattern. IS elements are highlighted in yellow.

Genomic comparison of *V. cholerae* strains reveal that the phage variation dynamics are of the “additive” kind, where small gene cassettes are integrated into the 3’ part of the gene coding for RtxA^60^. All the CTXΦ variants have conserved intergenic regions that separate and mark the insertion of these gene cassettes (*att* sites, marked as black arrows). These types of insertions have been reported previously in *V. cholerae* O139^61^. Interestingly, some strains such as 3528-08 (C17) can have up to five insertions (**Fig. 3C**). The first two of the five cassettes are identical, and each cassette has a copy of the two subunits of CT. The other three have lost the toxin genes and those coding for RstB, RstR and RstA, and acquired a replication endonuclease (**Fig. 3C**). The presence/absence of these cassettes is not correlated with the phylogeny and, when present, they have high identity indicating a fast turnover of these elements (**Fig. 3B**). Furthermore, these cassettes can be integrated into other genome locations, as in the case of the strain PivertUAT4Aug (**Fig. 3B**). In addition, using the sequence of *att* sites as reference we also found a collection of cassettes in the same order (including the one containing the two subunits of CT) in the close relative species *Vibrio mimicus*, suggesting another potential donor of CTXΦ to non-toxigenic *V. cholerae* (**Fig S3**).

#### 2. Vibrio Pathogenicity Island-1 (VPI-1)

The toxin coregulated pilus (TCP), an essential colonization factor, is encoded within VPI-1^31,32^. TCP mediates microcolony formation, which is crucial for intestinal colonization, and acts as the receptor of CTXΦ^33^. VPI-1 also encodes two critical virulence regulators: ToxT and TcpP^32,33^. VPI-1 can be transferred, via generalized transduction, between strains of *V. cholerae* and can also form circular intermediates^62,63^. The variable region of the island is located between the CDS VC_0821 (hypothetical protein) and VC_0846 (integrase) (**Fig. 4A**). The most conserved version of this region can be found in nine clades, four of them within C24 (2 from PCL) and five from C17 (**Fig. 4B**), including strain IRLE0081 isolated in this study (**Fig. 1C**). The average identity of the proteins was highly conserved compared to PCG (*ca*. >97%), however, we found some notable exceptions (**Fig. 4A**). First, among the proteins that are part of the TCP operon, we found high divergence rates in the master regulator ToxT (VC_0838) (**Fig. 4C**) and the toxin-coregulated pilus major pilin, TcpA (VC_0828) (**Fig. 4D**) with a median protein identity *ca*. 81% (**Fig. 4**). In addition, we found that the PCG allele of the colonization factor TcpF (VC_0837) (**Fig. 4E**) was only present in three strains (one PCL and two C17). In the rest, despite the conservation of synteny and similarity of the whole TCP operon, the identity of this protein was less than 40%, encoding a different allele of this poorly understood secreted protein^64–66^ (**Fig. 4B**). Finally, our genomic comparisons reveal that the gene coding for ToxR-activated gene A, TagA (VC_0820) was absent from the C17 genomes, while being retained in three of the four C24 genomes (**Fig. 4B**). Overall, despite widespread conservation in the synteny and size of the clusters of genes that comprise VPI-1 in the *V. cholerae* species, the existence of unique major allelic variations in a limited number of genes might act as a bottleneck in the emergence of PCG.

**Fig. 4.**
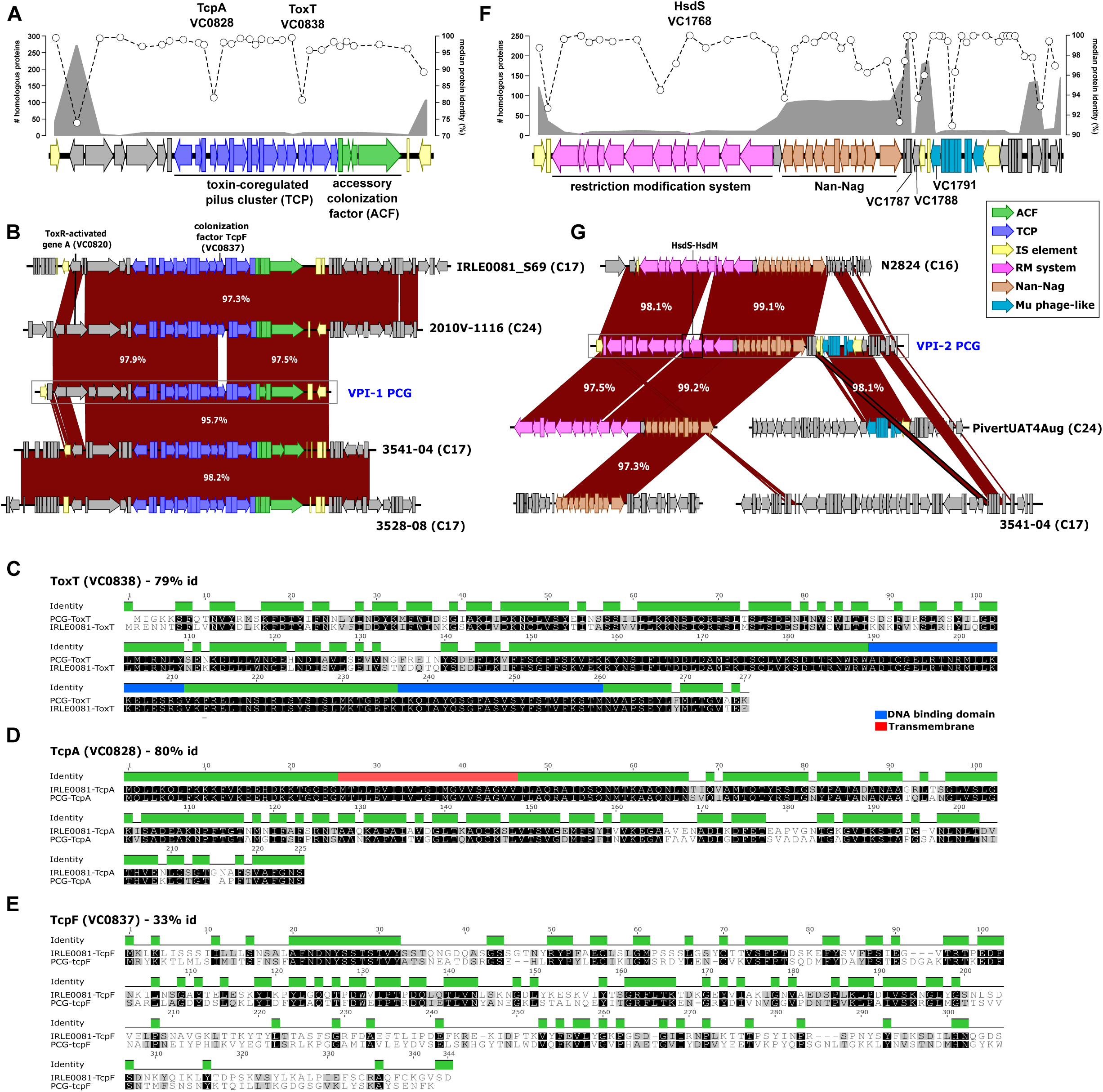
Modularity and allelic variations shape the evolutionary dynamics of Vibrio pathogenicity islands 1 and 2. Structures of **(A)** VPI-1 and **(F)** VPI-2. The area outlined in gray represents the number of homologous proteins found in the dereplicated *V. cholerae* genomes. The white circles with dotted lines indicate the median protein identity. **(B)** Genomic comparison of homologous VPI-1 in dereplicated *V. cholerae* genomes denote a reduced number of allelic variations in some PCG genes. Sequence alignment of the protein sequences of **(C)** ToxT **(D)** TcpA and **(E)** TcpF from PCG and environmental strain IRLE0081 (C17). **(G)** Genomic comparison of VPI-2 variants encoded in dereplicated *V. cholerae* genomes highlight pervasive modularity of the island. Clusters of genes with a known function follow the same color pattern. IS elements are highlighted in yellow.

#### 3. Vibrio Pathogenicity Island-2 (VPI-2)

VPI-2 is a large PAI that encodes the genes for sialic acid utilization^34^. The capacity to utilize sialic acid as a carbon and energy source confers *V. cholerae* a competitive advantage in the mucus-rich environment of the gut, where sialic acid availability is extensive^67^. Sialic acid catabolism also mediates a chemotactic response towards mucin and several environmental reservoirs of *V. cholerae*^68^. VPI-2 is excisable and form a circular intermediate^63,69^. Interestingly, during the excision event, there is crosstalk between VPI-2 and VPI-1^70^. Unlike other MGEs, the differential abundance of the gene clusters within VPI-2 suggests that the island is made up of different modules. Also, all but one of the IS elements of the island are widespread among several clades in diverse locations, which favours the transfer of the different modules. The canonical VPI-2 from O1 strains encodes three major modules, from 5’ to 3’: a) a restriction modification system (RM), b) the Nan-Nag cluster, and c) a Mu-phage like region (**Fig. 4F**). At the 3’ end of VPI-2 there is also a small group of genes encoding hypothetical proteins. The Nan-Nag cluster is present in 88 clades, whereas the distribution of the RM system is more limited, being encoded by only 7 clades (**Fig. 4F**). The Mu-phage like module is present in nine clades, however, there is no relationship between specific modules and species phylogeny. Like VPI-1, no clade other than PCG encode a PCG-like version of VPI-2. Furthermore, strains that contain the three individual modules sometimes encode them in different locations within the genome. For instance, strain PivertUAT4Aug (C24), the only one with the three major modules, encode the first two modules (Nan-Nag and RM) within a version of VPI-2, whereas the Mu-phage region is encoded in a different location of the genome (**Fig. 4G**). Similar to *rfb*T in the LPS gene cluster, VC_1788 and VC_1791 in this strain comprise two parts of a gene coding for the tape measure protein of phage Mu that have been truncated by the insertion of two transposases in the PCG, annotated as VC_1789 and VC_1790 (**Fig. 4F**). Although the synteny in the modules was conserved, and with high similarity (**Fig. 4G**), we found a high divergence in the VC_1768 gene coding for a restriction endonuclease S subunit (HsdS) between PivertUAT4Aug and the PCG-like version of VPI-2 with a protein identity of 64% (**Fig. 4G**). Another instance of this modular distribution is exemplified by strain N2824 (C16), which possesses the first two modules but not the Mu-phage. Furthermore, the strain has lost the genes encoding the HsdS and HsdM proteins from the RM cluster involved in DNA sequence recognition and methylation, respectively (**Fig. 4G**). Strain 3541-04 (C17) encodes a different cluster of genes at the insertion site of the Nan-Nag cluster and RM system, with the Nan-Nag cluster being inserted in another chromosomal region (**Fig. 4G**). Overall, VPI-2 in PCG must be understood based on its unique modular arrangement not by the individual units that constitute it and the possibility of allelic variability in *hsdS* playing a unique role in the pandemic group. Its four modules are associated with IS elements that are widely distributed among *V. cholerae* environmental strains and scattered throughout the genome. This leads to the convergence of all modules in the same genome being extremely low which contributes to the rarity of PCG.

#### 4. Vibrio Seventh Pandemic Island I (VSP-I)

The role of VSP-I in *V. cholerae* pathogenesis is not as clearly defined as other MGEs. VSP-I has been suggested to play an environmental role related to chemotaxis^71^ and encodes a regulator involved in intestinal colonization (VC_0177)^35^. We found homologs of the eleven genes that comprise this island in ten *V. cholerae* clades (**Fig. 5A**). Although four of them are found in strains isolated from this study, IRLE0062, IRLE0049, IRLE0042 and IRLE0077, that belonged to the same clade. The greatest sequence divergence corresponds to its integrase (VC_0183) and an XRE family transcriptional regulator (VC_0176) encoded within the island (**Fig. 5A**). Interestingly, only two strains encode a PCG-like VSP-I, V060002 (C24), and VcCHNf4 (C5). Surprisingly, V060002 encodes two identical copies of VSP-I in close proximity to each other with the LPS cluster located between them (**Fig. 5B**). The landscape of evolutionary possibilities such as modularity is much more limited for VSP-I than for the other MGEs due to its small size. It is also the rarest of all the islands with one transcriptional regulator, VC_0176, exhibiting high allelic diversity and very limited distribution. The rarity of the island and diversity of the regulator suggests that this is a critical bottleneck in the emergence of the Seventh Pandemic El Tor strains.

**Fig. 5.**
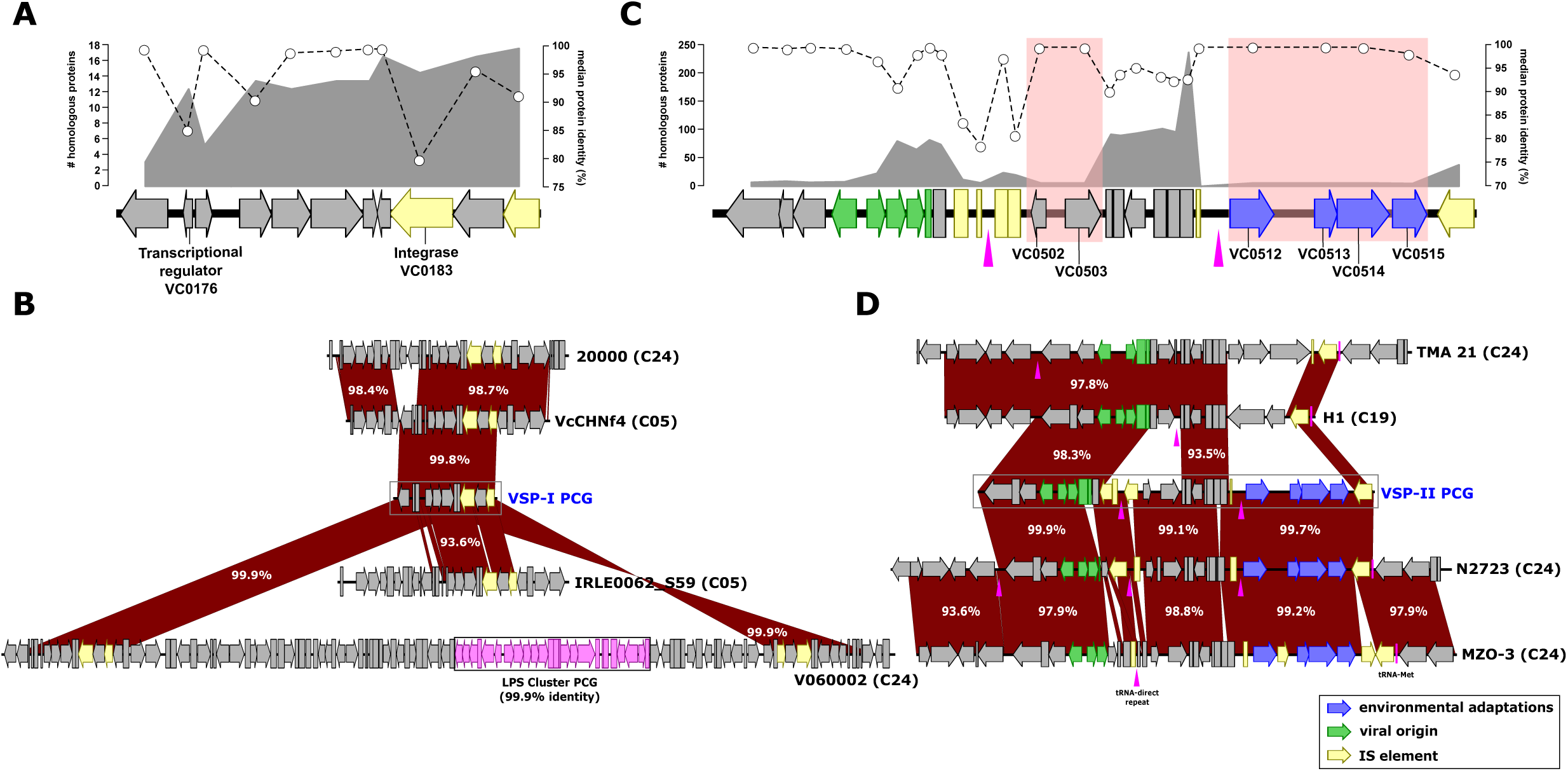
Distribution and evolutionary dynamics of the Vibrio seventh pandemic islands I and II. Structures of **(A)** VSP-I and **(C)** VSP-II. The gray area represents the number of homologous proteins found in the dereplicated *V. cholerae* genomes. The white circles with dotted lines indicate the median protein identity. Red rectangles on the genes indicate their exclusive presence in the PCL. Pink triangles indicate the presence of tRNA-direct repeat genomic islands. **(B)** Genomic comparison of homologous VSP-I across *V. cholerae* genomes. **(D)** Genomic comparison of VSP-II variants encoded in dereplicated *V. cholerae* genomes highlight pervasive modularity of the island. IS elements are highlighted in yellow.

#### 5. Vibrio Seventh Pandemic Island II (VSP-II)

The presence of VSP-II in clinical and environmental strains might be associated with environmental survival and fitness of the bacterium^36–38^. A group of *V. cholerae* strains that caused an outbreak in Florida associated with oyster consumption encoded a novel bacteriocin and a pyocin within the VSP-II element, proteins typically associated with bacterial competition^38^. Our analyses indicate that, similar to VPI-2, VSP-II is highly modular with several clusters of genes that have a tRNA-Met insertion point. The direct repetition of the end of the tRNA (highlighted as purple triangles) marks the insertion points of each module, which in most cases match the presence of IS elements (**Fig. 5C and 5D**). Overall, this island serves as another example of an “additive” island with three small modules. Module 1 comprises genes VC_0490 to VC_0498, five of which are of viral origin but lack essential proteins for packaging, indicating that it is a defective prophage. Module 2 encompasses VC_0502 to VC_0510, however, the lack of possible annotation of its genes precludes from assigning a specific function to this module. Nonetheless, VC_0502 and VC_0503 from this module are only present in strains from C24 (**Fig. 5C**). Module 3 is comprised of four genes (VC_0512 to VC_0515): VC_0512 and VC_0514 encode methyl-accepting chemotaxis proteins, VC_0513 a transcriptional regulator and VC_0515 a predicted signal transduction protein. These four genes are only present in C24 strains and according to their annotation could be associated with environmental adaptations (**Fig. 5D**). Interestingly, there are no identical versions of VSP-II outside of PCG. The closest versions are encoded by strains N2723, MZO-3, and V060002 belonging to C24, which are the only ones that possess a similar PCG-like modularity and synteny of VSP-II (**Fig. 5D**). N2723 has lost genes coding for two hypothetical proteins (VC_0496 and VC_0509) and a transcriptional regulator (VC_0497) and VSP-II from strain MZO-3 lost a transposase (VC_0501) in Module 1 and gained another in Module 3 (**Fig. 5D**). In the rest of *V. cholerae* strains, VSP-II only maintains some of these modules or clusters of genes, with the addition of other types of clusters being common. Finally, our analyses did not reveal any gene that exhibits high rates of divergence within the island, suggesting allelic diversity might not act as a major constraint in this element.

### Allelic variations confer a competitive advantage during intestinal colonization

We identified a set of genes encoded within the elements described above that are dispersed in the phylogeny of the species and possess unique variations in PCG. As stated above, we hypothesized that they might act as bottlenecks for the emergence of toxigenic clones of *V. cholerae* and shed light on the evolution of PCG. In order to test this, we constructed several isogenic mutants where we exchanged the wild-type PCG allele for an environmental allele as specified in **Table 1**. Specifically, we mutated the genes encoding:

**a) RfbT** (VC_0255). A FkbM family methyltransferase located in the LPS cluster associated with Inaba-to-Ogawa serotype conversion^54^. The PCG allele is truncated by transposases and was replaced by the allele of *V. cholerae* A325, which has the complete version of the gene.
**b) TcpA** (VC_0828). The TCP major pilin subunit is encoded within VPI-1 and has an average protein identity of 81%. We exchanged the PCG *tcpA* allele for the one encoded by the IRL isolate *V. cholerae* IRL0081.
**c) ToxT** (VC_0838). The master virulence regulator ToxT is also encoded within VPI-1 and has an average identity of 81%. We exchanged the PCG *toxT* allele for the one encoded by the IRL isolate *V. cholerae* IRL0081.
**d) TcpF** (VC_0837). The secreted factor TcpF is also encoded within VPI-1 with only three strains encoding the PCG-allele. The rest had an average identity less than 40%. We exchanged the PCG *tcpF* allele for the one encoded by the IRL isolate *V. cholerae* IRL0081.
**e) VC_1791.** A tape measure protein in the Mu phage-like module encoded within VPI-2. The PCG allele is truncated by the insertion of two transposases and was replaced by the allele of *V. cholerae* strain PivertUAT4Aug, which has the complete version of the gene.
**f) HsdS** (VC_1768). This restriction endonuclease S subunit encoded within VPI-2 has the lowest median identity of the entire island (64%). We exchanged the PCG *hsdS* allele for the one encoded by the *V. cholerae* strain PivertUAT4Aug.
**g) VC_0176.** An XRE family transcriptional regulator encoded within VSP-1 that has the lowest median identity of the entire island (85%). We exchanged the VC0176 allele from the IRL isolate *V. cholerae* IRL0062.

**Table 1.** List of strains used in this study.

| Strain | Description | Source |
| --- | --- | --- |
| *V. cholerae* C6706 | Wild-type, clinical *V. cholerae* O1 El Tor strain C6706; Sm<sup>R</sup> | Lab collection |
| *E. coli* S17 λ pir | Host for cloning vector | Lab collection |
| *ompU*<sup>GBE1114</sup> | *V. cholerae* C6706 encoding *ompU* allele from environmental strain *V. cholerae* GBE1114 | 22 |
| *rfbT*<sup>A325</sup> | *V. cholerae* C6706 encoding *rfbT* allele from environmental strain *V. cholerae* A325 | This study |
| *tcpA*<sup>IRLE0081</sup> | *V. cholerae* C6706 encoding *tcpA* allele from environmental strain *V. cholerae* IRLE0081 | This study |
| *toxT*<sup>IRLE0081</sup> | *V. cholerae* C6706 encoding *toxT* allele from environmental strain *V. cholerae* IRLE0081 | This study |
| *tcpF*<sup>IRLE0081</sup> | *V. cholerae* C6706 encoding *tcpF* allele from environmental strain *V. cholerae* IRLE0081 | This study |
| *hsdS*<sup>Pivert</sup> | *V. cholerae* C6706 encoding *hsdS* allele from environmental strain *V. cholerae* PivertUAT4Aug | This study |
| VC1791<sup>Pivert</sup> | *V. cholerae* C6706 encoding tape measure (VC1791) allele from environmental strain *V. cholerae* PivertUAT4Aug | This study |
| VC0176<sup>IRLE0062</sup> | *V. cholerae* C6706 encoding VC0176 allele from environmental strain *V. cholerae* IRLE0062 | This study |
| VC0176<sup>IRLE0062::</sup><br>*rfbT*<sup>A325</sup> | Double mutant; *rfbT*<sup>A325</sup> environmental allele introduced into the *V. cholerae* C6706 encoding VC0176<sup>IRLE0062</sup> | This study |
| Double mutant::<br>VC1791<sup>Pivert</sup> | Triple mutant; VC1791<sup>PivertUAT4Aug</sup> environmental allele introduced into the *V. cholerae* C6706 double mutant | This study |
| Triple mutant::*ompU*<sup>GBE1114</sup> | Quadruple mutant; *ompU*<sup>GBE1114</sup> environmental allele introduced into the *V. cholerae* C6706 triple mutant | This study |
| Quadruple mutant::*toxT*<sup>IRLE0081</sup> | Quintuple mutant; *toxT*<sup>IRLE0081</sup> environmental allele introduced into the *V. cholerae* C6706 quadruple mutant | This study |
| Quintuple mutant::*tcpA*<sup>IRLE0081</sup> | Sextuple mutant; *tcpA*<sup>IRLE0081</sup> environmental allele introduced into the *V. cholerae* C6706 quintuple mutant | This study |

We performed competitive assays using the infant mouse model of infection to determine whether the PCG-associated allelic variations of these genes confer a fitness advantage to toxigenic *V. cholerae*. Our results indicate that the PCG allele of *tcpF* is essential during the infectious process, with the mutant strain *tcpF^IRL^*^0081^ exhibiting loss of intestinal colonization, similar to the Δ*toxR* strain used as negative control, which is avirulent (**Fig. 6A**). On the other hand, the allelic specificity of HsdS does not appear to play a direct role in the emergence of PCG as the competitive index (CI) of the *hsdS^PIVERT^* mutant is close to 1 (**Fig. 6A**). The mutant strains for the other 5 genes exhibit a varying degree of fitness loss during intestinal colonization with *tcpA^IRL^*^0081^ showing the largest decrease (∼3-logs), followed by *toxT^IRL^*^0081^ (∼1-log), *VC1791^PIVERT^* (∼1-log), *rfbT^A3^*^25^ (∼7-fold), and *VC0176^IRL^*^0062^ (∼5-fold) (**Fig. 6A**).

**Fig. 6.**
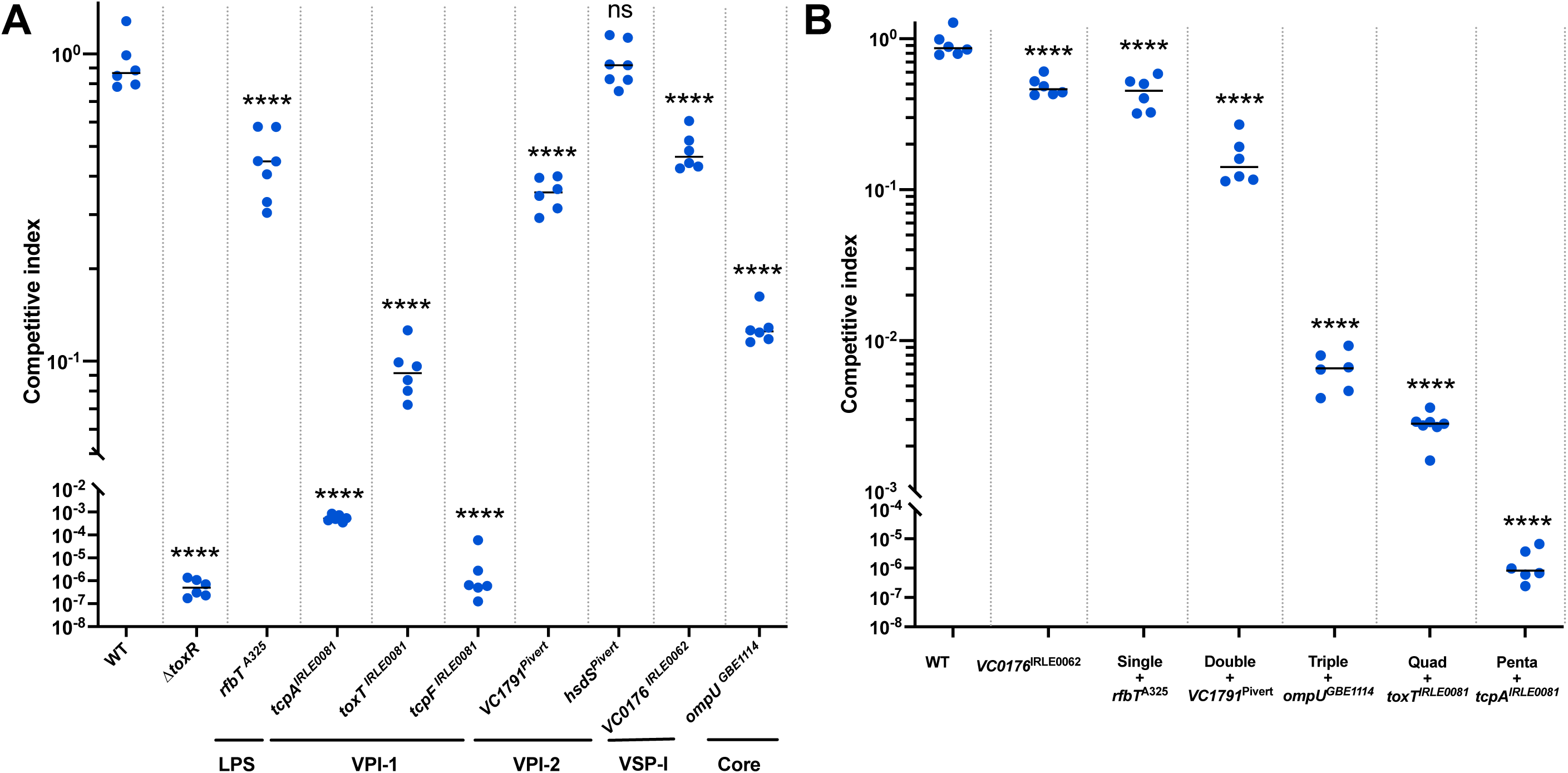
Allelic variations act as fitness bottleneck during intestinal colonization. Isogenic *V. cholerae* mutant strains were orogastrically inoculated in 3-5-day-old CD-1 mice to perform competition assays. Strains were monitored for colonization of the small intestine. The relative colonization data is represented as competitive indices (CI) in comparison to a co-infected *V. cholerae* C6706 Δ*lacZ* strain. **(A)** Isogenic strains encoding individual non-PCG alleles exhibit different competitive indices during intestinal colonization. **(B)** Combination mutants of the non-PCG alleles demonstrate an additive decrease in colonization efficiency. n = 6 per strain. *p*-values indicate significance compared to the WT strain, unless otherwise specified, and are calculated by 2way ANOVA in Šidák’s multiple comparisons test; *** <0.0001, **** < 0.00001.

We recently determined that allelic variations in core genes, in the form of virulence adaptive polymorphisms (VAPs), confer critical preadaptations for the emergence of pathogenic traits^22^. For instance, toxigenic strains encoding environmental alleles of the major outer membrane porin OmpU from the strain exhibit bile sensitivity and reduced resistance to host antimicrobials^22,7^^2^. Furthermore, those strains (e.g., *ompU^GBE1114^*) show a colonization defect in the infant mouse model (∼1-log) (**Fig. 6A**)^22^. In order to unveil the potential relationship among these allelic variations (MGEs and core genome), we constructed a set of mutant strains combining these alleles (**Fig. 6B**). We excluded those that showed no defect in colonization (*hsdS^PIVERT^*) or led to complete loss of colonization (*tcpF^IRL0081^*). We constructed the mutants on the background of *VC0176^IRL0062^*, as it was the one that showed the lowest effect in the CI (**Fig 6A**) and exchanged the alleles in ascending order based on their effect on the CI. Specifically, first we constructed a double mutant by introducing *rfbT^A325^* that exhibits a ∼5-fold decrease in its competitive index. Next, in the double mutant background, we constructed a triple mutant by introducing *VC1791^PIVERT^* (∼9-fold). Subsequently, we constructed a quadruple mutant by introducing *ompU^GBE1114^*(∼2.2-log) and a quintuple by introducing *toxT^IRL0081^* in the previous background (∼2.9-log). Finally, we generated a sextuple mutant encoding the previous 5 environmental alleles and *tcpA^IRL0081^* which is fully defective for intestinal colonization. Overall, there is a variable and non-linear effect between the alleles, with the largest effect being that of the introduction of *ompU^GBE1114^* and *tcpA^IRL0081^* (**Fig. 6B**). Phenotypic analyses indicate that the complete loss of colonization exhibited by *tcpF*^IRL0081^ and the sextuple mutant is not due to a growth defect, loss of motility, or changes in their ability to form biofilm (**Fig S4**). Our results demonstrate that allelic variations in MGEs, LPS cluster and core genome act as an additive yet stringent bottleneck that explicates the confined nature of toxigenic *V. cholerae* isolates and the PCG.

### PCG alleles do not influence colonization of model environmental reservoirs

Colonization of new niches or hosts is often associated with fitness trade-offs that make those adaptations beneficial within one context while detrimental on another^73,74^. In order to determine potential environmental fitness trade-offs associated with the PCG alleles, we examined the colonization dynamics of the two isogenic mutant strains that exhibit complete loss of intestinal colonization, *tcpF*^IRL0081^ and the sextuple, during colonization of three model environmental reservoirs: crustaceans, cyanobacteria and molluscs.

**a) Crustaceans**. We developed a model of infection using the brine shrimp *Artemia salina* to examine the interactions of *V. cholerae* with crustaceans (**Figs 7A, 7B and S5**). First, we examined the colonization dynamics of *V. cholerae* C6706 at different concentrations and time points (**Fig S5**). We determined that infection with 10^7^ CFU/mL for 48 hours leads to consistent colonization of *A. salina* nauplii (**Fig S5**). Surprisingly, even though we expected *V. cholerae* to colonize the chitinaceous surface of *A. salina*, the bacterium primarily colonizes the gut space of the nauplii (**Fig 7B**). Next, we performed co-infection studies using a similar approach as those in the infant mouse. Our competition assays demonstrate that mutant strains encoding *tcpF*^IRL0081^ or the sextuple mutant colonize *A. salina* at similar levels as the WT strain with a CI ∼1 (**Fig 7C**).
**b) Cyanobacteria**. We used *Microcystis aeruginosa* as a model system to investigate the interactions of *V. cholerae* with cyanobacteria as the bacterium has been found associated with this species between cholera outbreaks in endemic areas^68,75^. Competition assays demonstrate that, similar to *A. salina*, both mutants colonize the mucilaginous sheath of *M. aeruginosa* comparably to the WT strain.
**c) Mollusks**. In order to examine the interaction of *V. cholerae* with molluscs, we extracted the mucus-containing gills of Eastern oysters (*Crassostrea virginica*) and performed competition assays using their homogenates (see materials and methods). Similarly, the sextuple mutant and *tcpF*^IRL0081^ exhibit a CI in oyster gill homogenates analogous to the WT toxigenic *V. cholerae* isolates (**Fig 7C**). Overall, our results indicate that no significant fitness advantage or trade-off is associated with the acquisition of PCG alleles in these model systems.

**Fig 7.**
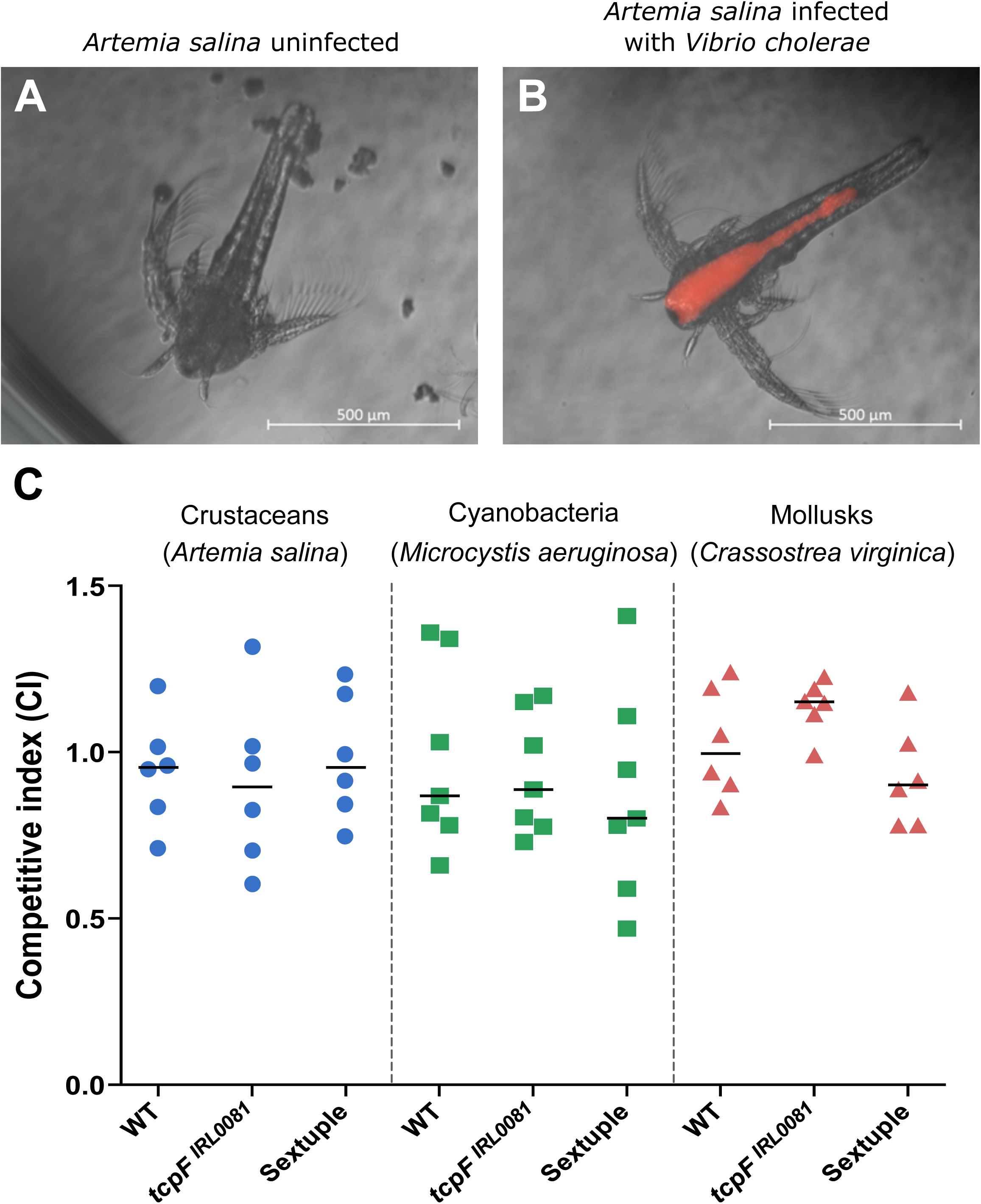
PCG alleles do not influence colonization of model environmental reservoirs. **(A)** Image of *Artemia salina* uninfected with *V. cholerae*. **(B)** RFP-tagged *V. cholerae* C6706 cells are ingested by *A. salina* colonizing and proliferating to fill the gut space. **(C)** Competitive indices of *tcpF*^IRL0081^ and the sextuple mutant in three model environmental reservoirs: Crustaceans (*A. salina*), cyanobacteria (*Microcystis aeruginosa*) and mollusks (gill homogenates of the Eastern oyster, *Crassostrea virginica*). The mutants do not exhibit any significant differences in CI during colonization of these reservoirs. n = 6. *p*-values indicate significance compared to the WT strain and are calculated by 2way ANOVA.

## CONCLUSIONS

In this study, we took a systematic approach towards unraveling the evolutionary history of the enigmatic Pandemic cholera group (PCG), which, to date, remains the only clade within the *V. cholerae* species capable of causing the severe diarrheal disease cholera. Phylogenomic analyses of the species reveal a cluster-based evolution of *V. cholerae* with several hierarchical levels leading to the emergence of the PCG. Our investigations on the abundance and distribution of the virulence-associated genes and MGEs suggest the existence of several layers in the evolutionary dynamics and emergence of *V. cholerae* (**Fig. 8**).

**Fig. 8.** Model of emergence of the Pandemic Cholera Group in *V. cholerae*. The evolutionary history of PCG suggests that the emergence of this group has not been a linear process and contains a series of major bottlenecks that explains its rarity. **(A)** A specific genomic background containing preadaptations to virulence in the core genome (red arrows) is essential for the emergence of virulence traits in *V. cholerae*. Green arrows: strains not encoding these preadaptations. **(B)** The acquisition of certain gene clusters is critical (e.g., CTXΦ, VPI-1, etc.) together with (**C)** the presence of specific modules within these gene clusters and **(D**) certain unique allelic variations within them. **(E)** The correct combination of all the elements of this complex mosaic provides the foundation for the emergence of pathogenic clones within an environmental population of *V. cholerae* (red arrow).

Our analyses indicate that **a)** strains from *V. cholerae* PCG have acquired the gene clusters encoding the major virulence genes in a modular fashion, **b)** those MGEs are more widespread than previously thought within the *V. cholerae* species, **c)** a select set of genes dispersed in the phylogeny of the species have a very limited distribution, and **d)** unique allelic variations in PCG act as stringent bottlenecks for emergence. Pangenome analyses of the *V. cholerae* species reveal that, besides the acquisition and modularity of the main MGEs, the absence or presence of other genes does not appear to play a major role in the emergence and evolution of the PCG (**Supp. text and Fig. S6A**). Furthermore, the phylogeny of *V. cholerae* shows that we can only trace the evolutionary history of the LPS cluster (**Supp. text and Fig. S6B**). This island-like element is exchanged by homologous recombination and appears to have slower replacement dynamics than the MGEs. On the other hand, the MGEs are additive genomic islands composed of several modules bordered by IS elements. These modules have a rapid turnover and remain associated with each clade for a short time. In addition, these modules have the potential to be inserted into other regions of the genome. This mosaic-like degree of molecular sophistication explains the absence of a complete set of PCG-like elements in any other *V. cholerae* clade. Indicating that, in order to analyze the evolutionary history of these mobile elements, it is necessary to examine the specific modules and individual alleles of some genes within them rather than investigating presence or absence of the elements (**Fig. 8**).

To examine whether these intra-modular allelic variations act as an evolutionary constraint that govern the emergence of toxigenic *V. cholerae*, we constructed isogenic mutant strains where we exchanged the PCG-associated allele for an environmental one and investigated their fitness during intestinal colonization. We determined that the PCG-associated alleles lead to a competitive advantage during intestinal colonization. Surprisingly, we uncovered a differential effect between alleles of genes encoded within MGEs, the LPS, and core genome. Specifically, we found that some PCG-associated alleles are essential for successful colonization, such as *tcpF*, whereas others provide non-linear competitive advantages during infection that ultimately act as a critical bottleneck that clarifies the isolated emergence of PCG. Future research will elucidate the specific role and relative importance of these alleles and their potential interactions. Elucidating this will open novel research avenues and provide essential insights into the factors and network of genes associated with the emergence of virulence traits in bacteria. Finally, it appears that none of the three environmental reservoirs we examined in this study exert the ecological pressures associated with the selection of the PCG alleles as the mutants exhibit a similar competitive index as the wild-type strain. Given that these variations originate from the environment it is possible that a combination of abiotic and biotic factors might select for the PCG alleles in the aquatic ecosystem. Uncovering the environmental drivers fostering the selection of factors associated with pathogen emergence will provide critical insights towards the development of surveillance networks. Overall, our investigations provide unique insights and perspectives to elucidate the barriers associated with the emergence of human pathogens and can be extrapolated to the study of niche colonization and other complex phenomena.

## MATERIALS AND METHODS

### Sampling sites

We collected samples at two environmentally distinctive locations along the IRL (Eastern Florida, USA) in three sampling events. For each sampling campaign, we obtained samples from several fractions: three associated with the water bodies (water filtered through 20μm, 5μm, and 0.22μm) and one from the sediment. One of the sampling locations, Fellers House Field Station (N28°54’25.315”; W80°49’15.017”), is located within the federally-protected Canaveral National Seashore. The second sampling site, Shepard Park, is located in the Port St. Lucie area (N27°11’48.864”; W80°15’33.172”) that due to urbanization and agricultural expansion, experiences nutrient over-enrichment leading to excessive macroalgal bloom (**Fig. 1A**)^45^.

### Isolation of *V. cholerae*

**a).** *Water*. Isolation of *V. cholerae* from water samples collected at the sampling locations was performed using a modified protocol from Huq *et al* ^76^. Briefly, water samples (∼3L) were collected in triplicate from each site and stored cold until arrival at the laboratory. To investigate the prevalence of *V. cholerae* in various fractions of the aquatic column and to separate particle-associated and free-living bacteria, 500 mL of collected water was filtered successively through 20µm, 5µm and 0.2µm filters (Sterlitech) using a vacuum filtration system. The respective membrane filters harboring potential *vibrio* spp. were then suspended in 25ml phosphate buffered saline (PBS) and vortexed vigorously to release the contents. Each resuspended sample was then inoculated (1:10 v/v) in alkaline peptone water (APW) and incubated at 37°C for 12-14 hrs. This was repeated for each replicate. Enriched cultures were subjected to growth on a series of selective agar specific for the isolation of *V. cholerae.* First, enriched cultures were serially diluted 10-fold and plated on CHROMagar Vibrio (CaV; CHROMagar), a chromogenic media on which *V. cholerae* forms turquoise blue colonies. Subsequently, turquoise blue colonies from CaV were transferred onto Thiosulfate Citrate Bile Salts Sucrose (TCBS; Sigma) agar plates. Colonies that appeared turquoise blue on CaV followed by yellow on TCBS were preliminarily identified to be *V. cholerae*. **b)** *<u>Sediment</u>.* Isolation of *V. cholerae* was done using established methods^45^. Briefly, sediment samples were collected in triplicates from each sampling site using universal corer. 25 g of the sediment samples from each replicate was weighed and suspended in 25 ml of PBS (Gibco). Samples were homogenized, enriched, and studied for the presence of *V. cholerae* as described above. **c)** *<u>Cyanobacteria</u>.* Cyanobacteria floccules were pelleted, supernatant removed and fresh APW added to homogenize the sample. Samples were enriched and studied for the presence of *V. cholerae* as described above.

### Strains and growth conditions

*V. cholerae* strains were routinely grown on Luria Bertani (LB) agar at 37 °C for approximately 16hrs, unless otherwise stated. For routine liquid cultures, isolated colonies selected from agar plates were grown aerobically at 37°C in LB broth for ∼16hrs. Tryptone broth (tryptone 10 g/L, NaCl 5 g/L), artificial sea water and M9 minimal media (Fisher) supplemented with 0.1% glycerol were used for biofilm formation, studies using *Artemia* and oyster gill homogenates, respectively. Growth media was supplemented with the following antibiotics and reagents, as needed: polymyxin B 50 U/mL, kanamycin 45 µg/mL, streptomycin 1000 µg/mL, gentamycin 15 µg/mL, X-gal 40 µg/mL.

### Clonal analysis of *V. cholerae* isolates

To confirm isolates of *V. cholerae,* presence of the species-specific gene *ompW* was determined by PCR^46^. Genomic DNA from the isolates was extracted using the Gentra Puregene Yeast/Bact Kit (Qiagen) and used as template for PCRs. To identify the clonal populations of *V.* cholerae in the confirmed dataset, DNA fingerprinting patterns were examined using a combination of *Vibrio cholerae* Repeats-PCR (VCR-PCR) and Enterobacterial Repetitive Intergenic Consensus Sequence (ERIC)-PCR as described by Teh *et al* ^47^. Genomic DNA was used to perform VCR- and ERIC-PCR and resulting PCR products were electrophoresed and imaged using UVP ChemStudio (AnalytikJena) to observe fingerprinting patterns. Cluster analysis of DNA patterns was performed using BioNumerics v8.0 (Applied Maths, Inc.) and dendrogram generated based on the Unweighted Pair Group Method with the Arithmetic Mean (UPGMA) method^47,77,78^ (**Fig. S1).** Based on resulting DNA fingerprinting patterns, a total of 35 independent clones were identified and selected for whole genome sequencing (WGS) (**Table S1**). Genomic DNA of each clone was submitted to Microbial Genome Sequencing Center (MiGS) for WGS using the Illumina NextSeq 2000 platform.

### Genome assembly, gene prediction, and annotation

Reads were trimmed using Trimmomatic v0.36^79^ and assembled de novo with SPAdes version 3.11.1^80^. Coding DNA sequences (CDS) from the assembled contigs were predicted using Prodigal version 2.6.3 using *-a output.proteins -d output.genes -c -p meta* parameters^81^. Then, tRNAs were obtained using tRNAscan SE version 1.4^82^ together with ssu align version 0.1.1, and rRNA genes using meta rna^83^. Predicted proteins were compared against the National Center for Biotechnology Information nonredundant database (NCBI nr) using DIAMOND^84^. In addition, for functional annotation we used HMMscan version 3.1b2^85^ for the comparison against COG v2003 (update 2014)^86^ and TIGFRAM v15.0 (September 2014)^87^ databases.

### Recovery of *V. cholerae* genomes

A total of 1,127 *V. cholerae* genomes were downloaded from RefSeq (accessed July 2021) and subjected to a dereplication step with drep^88^ at 99 % nucleotide identity. Briefly, we force dRep to sort and select as seeds for the clustering genomes with better score. For instance, chromosomes in a single contig, followed by higher N50 and L50 values, and lastly by the completeness and contamination values calculated with checkm^89^. This resulted in 227 dereplicated reference *V. cholerae* genomes.

### Phylogeny of *V. cholerae* dereplicated genomes

Single nucleotide polymorphisms (SNPs) among *V. cholerae* dereplicated genomes and the PCG reference genome *Vibrio cholerae* El Tor N16961 (RefSeq accession number GCF_900205735.1) were calculated with PhaME^90^ with default options, resulting in a core genome length of 2,468,623 bp and 458,798 polymorphic positions considered. Then, a maximum likelihood phylogenetic tree of SNPs was constructed using iqtree v 1.6.12^91^ using the ultrafast bootstrap approach (5000 replicates) and the best fitted model GTR+F+R4.

### Abundance and distribution of pathogenicity-related islands

To determine virulence-associated mobile genetic elements and LPS cluster distribution and degree of prevalence of proteins within the studied *V. cholerae* genomes, we extracted these genomic regions from PCG reference genome (*V. cholerae* El Tor N16961) based on the manual annotation found in the NCBI Genbank. Then, proteins were extracted and used as a reference in a BLASTP search against the proteins from the rest of the *V. cholerae* species genomes^92,93^. We kept all proteins that matched against the pathogenic protein database with at least 70% identity and the alignment covered at least 70% of both proteins.

### Construction of mutant strains

*V. cholerae* constructs were all generated in the El Tor strain C6706 background following previously published methods^22^. Briefly, the environmental alleles of interest were synthesized commercially (Gene Universal) including 500 bp flanking regions from *V. cholerae* C6706 to facilitate recombination. The synthesized products were digested and ligated into the *V. cholerae* suicide vector pKAS154, electroporated into *Escherichia coli* S17λ-*pir* and selected on kanamycin plates. Following sequence verification, allelic exchange was carried out in *V. cholerae* C6706 using the appropriate antibiotic selection, as described^94^. Positive clones were first screened by colony PCR using flanking primers and confirmed by sequencing the exchange locus from purified genomic DNA. Fluorescent tagging of the strains was performed using the mini-Tn7 system that integrates at the *att* site between the *glmS* (*VC0487*) and *VC0488* genes in *V. cholerae*^95^. Briefly, triparental mating was performed using LB-grown overnight cultures of the donor *E. coli* S17λ-*pir* cells harboring pUC18T-mini-Tn7T-GFP or RFP expressed from a strong constitutive promoter, with the recipient *V. cholerae* cells and helper *E. coli* encoding the transposase gene on a suicide vector. The mating mixture were selected twice on LB-Pb-Gm plates and followed by two rounds of selection on LB-Sm plates, and subsequently confirmed by sequencing the Tn7 insertion site and fluorescence microscopy.

### Infant mouse colonization assays

Intestinal colonization competition assays in three-to-five-day-old infant mice were performed with LB-grown *lacZ*^+^ cells of the WT and allelic exchange mutants competed against an isogenic WT strain harboring a *lacZ* deletion, essentially as described^22,96,97^. The colonization efficiencies of the different strains in the small intestine harvested post infection are represented as competitive indices. Briefly, strains were inoculated from fresh LB plates into LB-Sm broth, incubated for 12 hours at 37 °C with aeration, and diluted 1:1000 fold in 1x PBS (pH 7.4, Gibco). The test (*lacZ*^+^) and control (*lacZ*^-^) strains were mixed 1:1 to obtain ∼10^6^ CFU/mL. To monitor successful inoculation, Evan’s blue dye (0.01% final, Sigma) was added to the cell mixture. Fifty microliters of the bacterial mixture, corresponding to ∼10^5^ CFUs, were then used to intragastrically inoculate 3-5-day-old infant mice and maintained at 30 °C post-infection. The exact input CFU numbers were determined by plating dilutions of the inoculum on LB-Sm-X-Gal plates. Approximately 22 hours post infection, the mice were euthanized, their small intestines harvested and transferred into 4 mL LB-10% glycerol stored on ice. The samples were then processed for CFU determination by homogenizing the tissues, serially diluting in 1x PBS (pH 7.4) and plating appropriate dilutions on LB-Sm-X-Gal plates. *In vitro* competition experiments were performed in parallel by inoculating 50 µL of the input CFU in LB-Sm broth and incubating for 22 hours at 37 °C with aeration and enumerating CFUs as above. Results are presented as the competitive index (CI), which is the ratio of the intestinal *lacZ*^+^ CFU (blue colonies) to *lacZ*^-^ CFU (white colonies) normalized to the input CFU ratio and *in vitro* competition CFU ratio.

### Colonization of environmental hosts

Bacterial inocula for infections of the environmental hosts were prepared essentially as described for the mice colonization assays but used at different dilutions and without the Evan’s blue dye. Coinfections were performed with GFP-tagged *lacZ-*positive test strains competing against RFP-tagged wild type strain harboring a *lacZ* deletion. **a)** *<u>Crustaceans</u>*. We modified previously established protocols as follows^98^, eggs of *A. salina* were hatched in artificial sea water supplemented with 2% sodium chloride (ASW), incubated at room temperature and aeration under a 12-hour light-dark cycle for 48 hours. Infections were performed in 200 µL ASW containing 20-30 individual crustaceans and the 10^7^ CFU/mL of bacteria. After 48 hours incubation at room temperature, samples were washed twice in ASW to remove unattached bacteria and either homogenized for determining CIs (as above) or used directly for microscopy. **b)** *<u>Cyanobacteria</u>*. We modified previously established protocols as follows^99^, *Microcystis aeruginosa* (UTEX LB 2385) were incubated in flasks containing BG-11 media (Gibco) under a 12-hour light-dark cycle with aeration (100 rpm) for about 14 days. *M. aeruginosa* cells were concentrated by centrifugation at 7500 rpm for 10 minutes, resuspended in BG-11 and co-cultured with 10^7^ CFU/mL *V. cholerae* cells at a final OD700 of 0.3-0.4 in a 200 µL reaction volume. Samples were harvested at 24 hours and CIs were determined on LB-Sm-X-Gal plates, as above. **c)** *<u>Mollusks</u>*. Mucus-containing gills of the Eastern oyster *Crassostrea virginica* were harvested in 1x PBS pH 7.4 (1 g/mL), homogenized and autoclaved^68^. M9 minimal media was supplemented with 0.1% sterile gill homogenate and inoculated with 50 µL of 10^7^ CFU/mL of bacterial inoculum. Subsequently, cultures were incubated for 24 hours at 37 °C under aerated conditions. CIs were determined by plating appropriate dilutions on LB-Sm-X-Gal plates.

### Phenotypic assays

**a)** <u>Growth on LB</u>. Overnight cultures were diluted to ∼10^5^ CFU/ml in fresh LB and 200 µL aliquots in flat-bottom 96-well plates (CytoOne) were monitored at OD595 nm, 37 °C for 24 hours in a Tecan Sunrise plate reader. **b)** <u>Biofilm formation</u>. These assays were performed as previously described with minor modifications^100^. Briefly, overnight cultures in tryptone broth were diluted 1:500 in fresh broth and 200 µL aliquots in 96-well polystyrene plates (CytoOne) were incubated at 37 °C for 24 hours. After washing to remove unattached cells, biofilm cells were stained with 0.1% crystal violet and quantified by measuring OD595 after elution with 50% glacial acetic acid. **c)** <u>Motility</u>. These assays were performed in soft agar plates (LB-0.3% agar) as previously described^68^. Diameter was measured for 12 hours every two hours.

### Pangenome analysis

To evaluate the pangenome of *V. cholerae*, in a first approach dereplicated sequences were grouped in clusters at 98% identity using cd-hit^101^, resulting in 34 clusters, four of them (C5, C11, C16, and C24) comprising 188 out of 262 genomes. Then, pangenomes were calculated for each cluster individually using PPanGGOLiN^102^ with default parameters. A second pangenome step with the protein families of clusters C5, C11 y C16 was performed using PPanGGOLiN to simplify the comparison among the PCG reference genome (*V. cholerae* El Tor N16961), C24 and the new superpangenome C5-C11-C16. Ortholog proteins among the three groups (El Tor, C24 and C5-C11-C16 were calculated using orthoMCL^103^ (> 30 % amino acid identity, > 70 % alignment). A Venn diagram showing proteins shared and not shared among groups was performed in R using the ggVennDiagram package (https://github.com/gaospecial/ggVennDiagram).

### Determination of Gain and Loss of genes in the pandemic cholera lineage (PCL)

In order to understand the flow of genes among pandemic *V. cholerae* and the closest genomes from cluster C24, those belonging to the PCL, gene presence and absence was tested using the gain loss mapping engine GLOOME^104^. The expectations and probabilities of both gain and loss events were estimated using stochastic mapping. The phyletic pattern of presence/absence of orthologous genes was then classified using hmmscan^85^ against the Pfam database v35.0^105^, using an evalue of < 1e^-5^ as a threshold for protein annotation.

### Microdiversity analysis

To identify which genes in each genome were under positive selection, we calculated the nonsynonymous (d*N*) and synonymous (d*S*) substitutions between ortholog proteins found in both the studied *V. cholerae* genomes and the reference PCG strain, we used the orthologr package in R^49^, a pipeline that identifies orthologous protein pairs, performs codon alignment and finally computes the final dN/dS values. We considered a gene under positive selection if the dN/dS ratio > 1.

## Data availability

The genomes have been deposited as BioSample SAMN38476510 to SAMN38476544 under BioProject PRJNA1035537.

## ACKNOWLEDGEMENTS

We thank Zac Carver for his technical assistance. This work was supported by a National Science Foundation (NSF) CAREER award (#2045671) and a Burroughs Wellcome Fund Investigator in the Pathogenesis of Infectious Disease award (#1021977) to SAM. MLP was supported by a “FLEX3GEN” PID2020-118052GB-I00 (cofounded with FEDER funds) from the Spanish Ministerio de Economía, Industria y Competitividad.

## AUTHOR CONTRIBUTIONS

SAM conceived the study. DB and TAG collected samples, isolated *V. cholerae* and sequenced genomes. MLP, AZS and JHM performed bioinformatic analyses. DB and CC constructed mutants and performed host colonization assays. The manuscript was written by MLP, DB and SAM. All authors read and approved the final version.

## SUPPLEMENTARY MATERIAL

**Table S1.** List of *Vibrio cholerae* collected in this study, together with genomic features and metadata.

**Table S2.** List of dereplicated (ANI>99%) *Vibrio cholerae* reference strains from the NCBI

**Table S3.** List of dereplicated (ANI>98%) *Vibrio cholerae* reference including IRL strains

**Table S4**. Shared family gene clusters in the pangenomes of the three hierarchical levels

**Table S5**. Summary of the gene gain/loss of the branching event 3 from the pandemic cholera lineage.

**Table S6**. Genes of the PCG under positive selection (dN/dS >1) compared to more than 60 non-clonal *V. cholerae* clades.

**Figure S1. Dendrogram representing the relatedness of *V. cholerae* isolates**. The dendrogram is based on PCR fingerprints generated by VCR-PCR. Groups of similarity were established using the UPGMA method in BioNumerics v8.0. The red circles indicate clonal isolates that were selected for whole genome sequencing.

**Figure S2. Pairwise comparison among the *V. cholerae* genomes.** The pairwise comparison was obtained using the average nucleotide identity (ANI) from strains isolated in this study and the Pandemic Cholera Group (*V. cholerae* El Tor N16961).

**Figure S3. Genomic comparison of CTX**Φ **phage**. Comparison between the CTXΦ phage genomic island encoded by Pandemic Cholera Group strains (*V. cholerae* El Tor N16961) and *Vibrio mimicus* strain 2011V-1073.

**Fig S4. Phenotypes of the *tcpF* ^IRL0081^ and sextuple mutants**. Isogenic *V. cholerae* C6706 strains encdoing either the *tcpF* ^IRL0081^ allele or the sextuple alleles were monitored for differences in **(A)** growth patterns in LB broth, **(B)** motility in 0.3% agar and **(C)** biofilm formation. n=6. *p*-values were determined using one-way ANOVA.

**Fig. S5. Artemia model of *V. cholerae* infection. (A)** CFU/ml of *V. cholerae* C6706 recovered post-infection at different time points and inoculum concentrations normalized to the input. Infection threshold (orange line) was determined by visualizing colonization using fluorescent microscopy. **(B)** RFP-tagged *V. cholerae* C6706 cannot colonize *A. salina* with an inoculum of 10^6^ CFU/ml after 48 hours and fluorescence cannot be detected. **(C)** Inocula of 10^7^ CFU/ml lead to consistent colonization of *A. salina* by RFP-tagged *V. cholerae* C6706 after 48 hours and red fluorescence can be detected.

**Fig. S6. Evolutionary trajectories and genome dynamics of *V. cholerae* PCG. (A)** *Pangenome analysis*. The Venn diagram shows the numbers of unique and shared gene families between the pandemic cholera group (purple), pandemic cholera lineage (blue), Cluster 24 (light blue) and the rest of clades (grey). **(B)** *PCL evolution*. The tree designates the evolutionary relationships of the different clades within the PCL highlighting the loss and gain of genes through the lineage. Black numbers indicate an overall gain and grey an overall loss. The colored circles represent the specific element as well as the percentage of completeness in each of the clades in reference to PCG. LPS, O1 lipopolysaccharide cluster; VPI-1, Vibrio pathogenicity island-1; VPI-2, Vibrio pathogenicity island-2; VSP-1, Vibrio seventh pandemic island-1; VSP-2, Vibrio seventh pandemic island-2. **(C)** *Microdiversity analysis*. PCG genes under positive selection compared to the rest of the clades. X-axis indicates the number of clades in which d*N*/d*S*>1 and the Y-axis shows the average of the d*N*/d*S* value.

