## Supplementary figures and images for "Allelic variations and gene cluster modularity act as non-linear bottlenecks for cholera emergence"

### Supplemental Figure 1

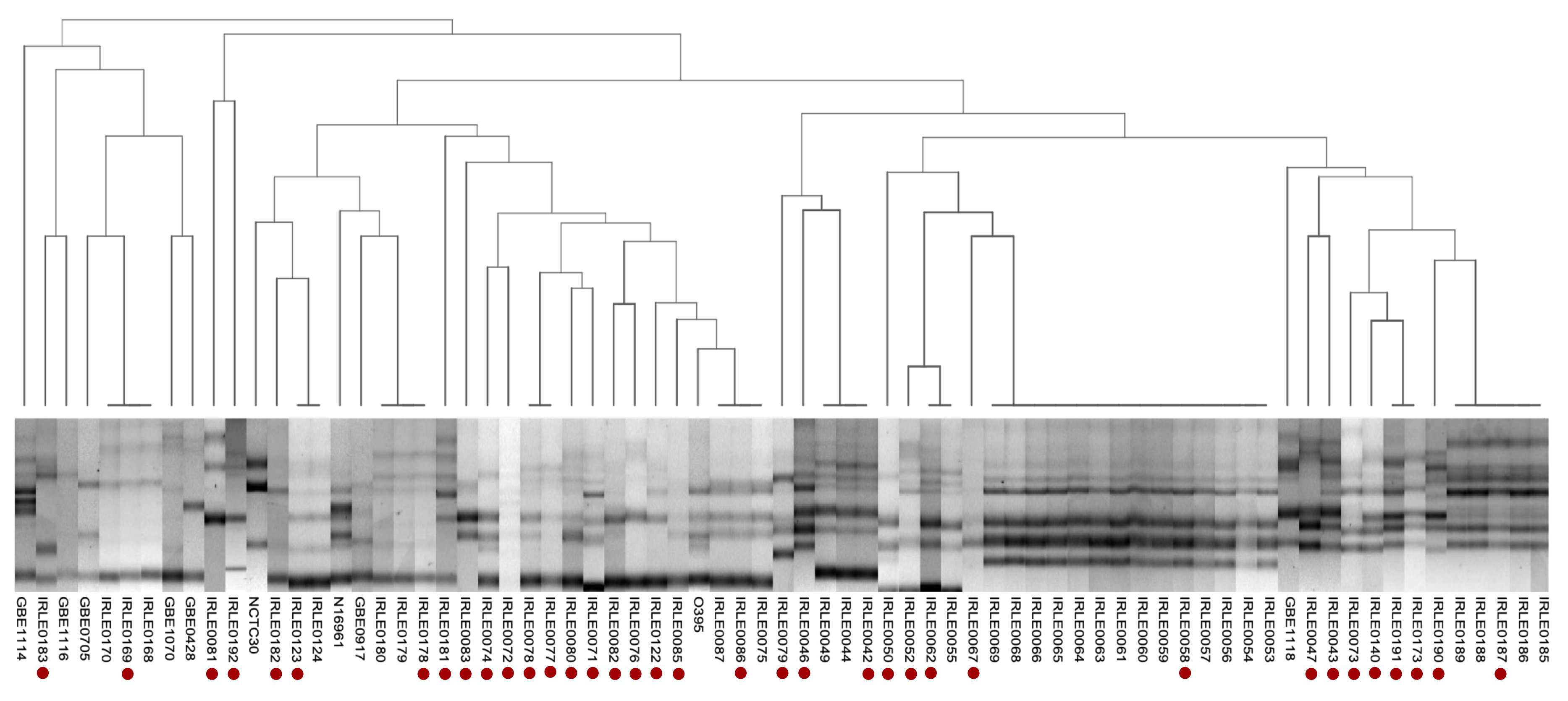

### Supplemental Figure 2

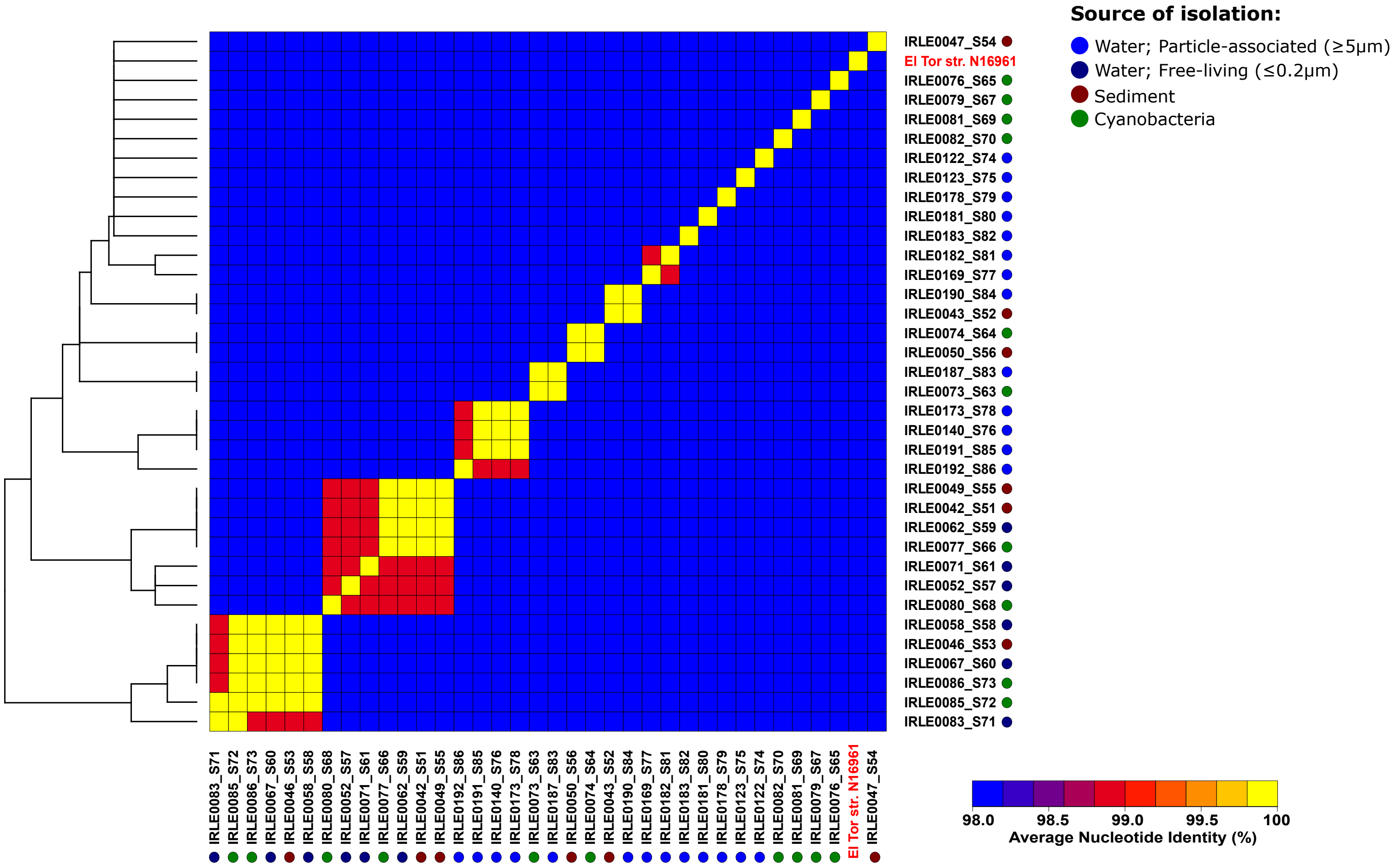

### Supplemental Figure 3

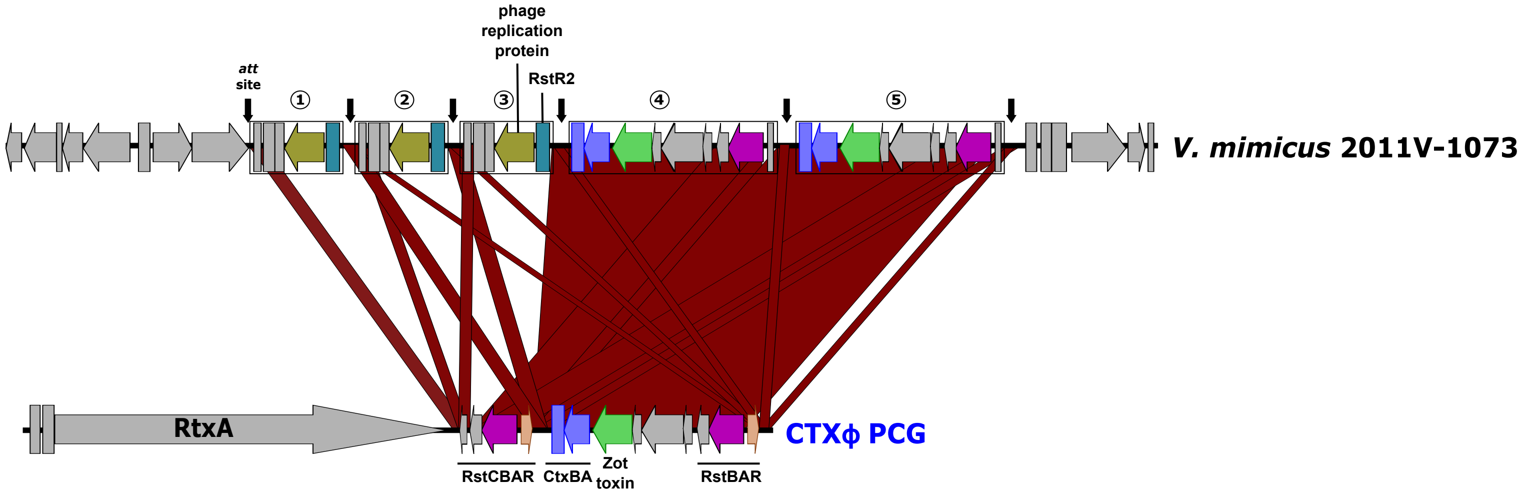
